# Highly Pathogenic Avian Influenza A (H5N1) Suspected in penguins and shags on the Antarctic Peninsula and West Antarctic Coast

**DOI:** 10.1101/2024.03.16.585360

**Authors:** Fabiola León, Céline Le Bohec, Eduardo J. Pizarro, Loïcka Baille, Robin Cristofari, Aymeric Houstin, Daniel P. Zitterbart, Gonzalo Barriga, Elie Poulin, Juliana A. Vianna

## Abstract

Suspected cases of Highly Pathogenic Avian Influenza (H5N1) were detected in Adélie penguins and Antarctic shags at the southernmost latitude so far in Antarctica, at two breeding sites out of 13 visited, using highly specific PCR assay. These first records mark the progression of the H5N1 panzootic into Antarctica.

## Introduction

The HPAI H5N1 clade 2.3.4.4b rapidly spread across continents within a two-year period starting from 2020, facilitated primarily by wild bird movements (Alkie et al., 2022). As the intensity of H5N1 clade 2.3.4.4b HPAI outbreaks continues to rise, the impact on wildlife has become significantly more severe (Charostad et al., 2023). The recent panzootic poses a significant conservation challenge, particularly on the health and survival of Antarctic wildlife. The challenge is exacerbated by the limited surveillance of Antarctic regions, complicating human monitoring and preventive measures. Moreover, colonial nesting seabirds are particularly vulnerable to disease transmission (Boulinier, 2023). These birds, whose reproductive behaviors involve nesting closely together for prolonged periods, face heightened risks of disease spread (Dewar et al., 2023). These circumstances significantly increase the threat to their health and population stability.

Although other strains of H5 avian influenza virus have been reported in seabirds in Antarctica before (Hurt et al., 2014; Barriga et al., 2016; Ogrzewalska et al., 2022), the pathogenic H5N1 has never been reported up to 2023. In October 2023, a sudden and marked increase in strandings and mortality was noted among skuas, kelp gulls, elephant seals, gentoo penguins and fur seals in Falkland/Malvinas, Bird Island, South Georgia, and the South Shetland Islands (Bennison et al., 2023), raising significant concern. Suspected mortalities of skuas were observed in Heroina Island, and Esperanza bay in Antarctica. More recently, in February 2024, positive cases were detected among Skuas in the Western Antarctic Peninsula (https://scar.org/library-data/avian-flu).

We conducted epidemiological surveys of seabird nesting sites in the Antarctic Peninsula, Weddell Sea, and the Antarctic western coast (Bellinghausen, Amundsen, and Ross Seas) in December 2023 and January 2024, and report our findings here.

## Methods

From December 2023 to January 2024, 13 seabird breeding sites were visited along the East and West coasts of the Antarctic Peninsula and the Antarctic western coast (Figure 1), on board Le Commandant Charcot PONANT vessel. A total of 115 birds from 4 species were captured, sampled, and released (see Table 1 for species and sample sizes). The procedure was conducted taking into account the recommendations provided by Biological Risk Assessment of Highly Pathogenic Avian Influenza in the Southern Ocean of SCAR (Dewar et al., 2023). The cloacal sampling protocol was carried out using 45 cm swabs. Samples were preserved in denaturant Viral Transport Medium VTM and refrigerated at 4°C. They were processed within the weeks following sampling.

**Table 1.**
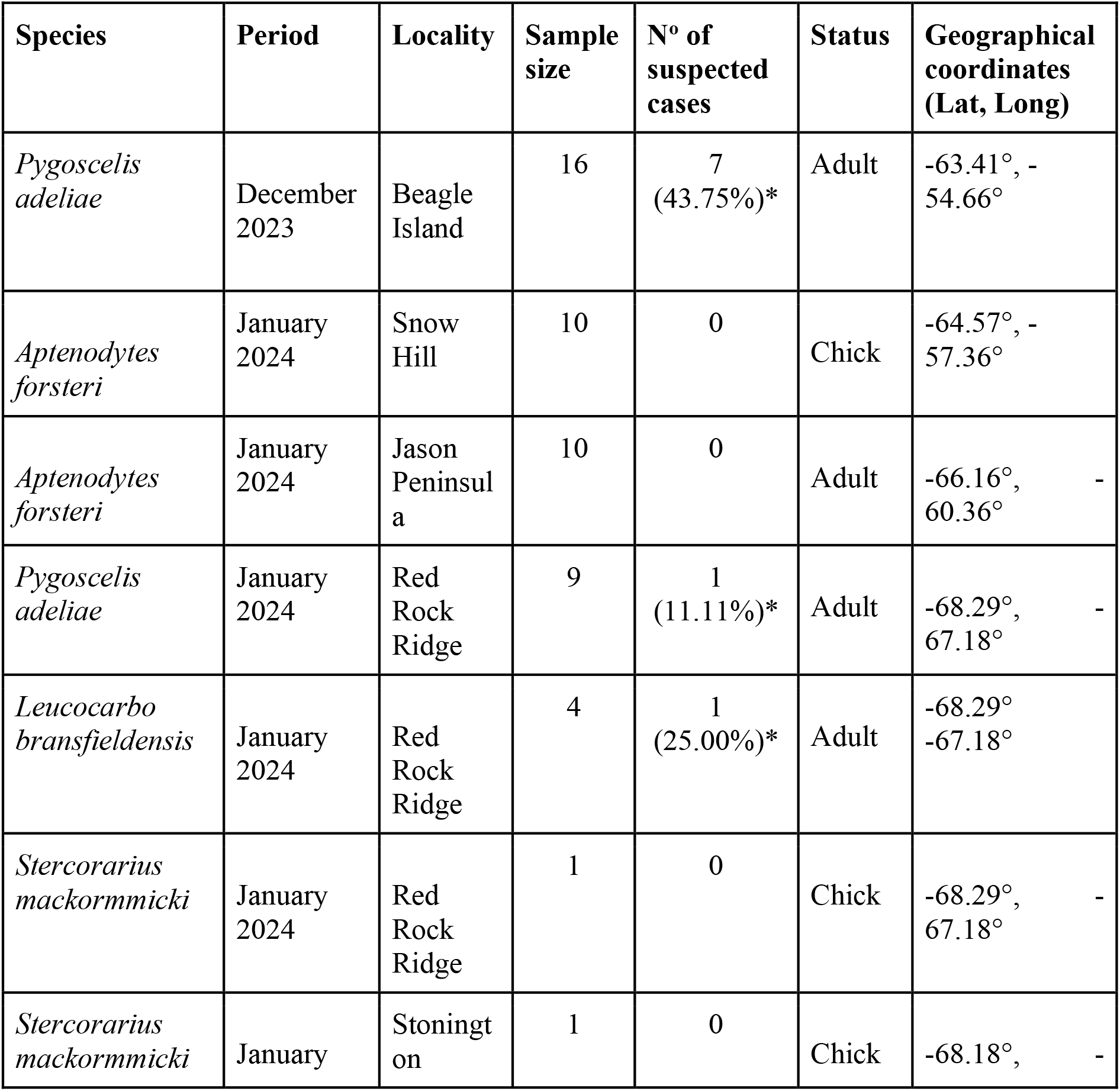
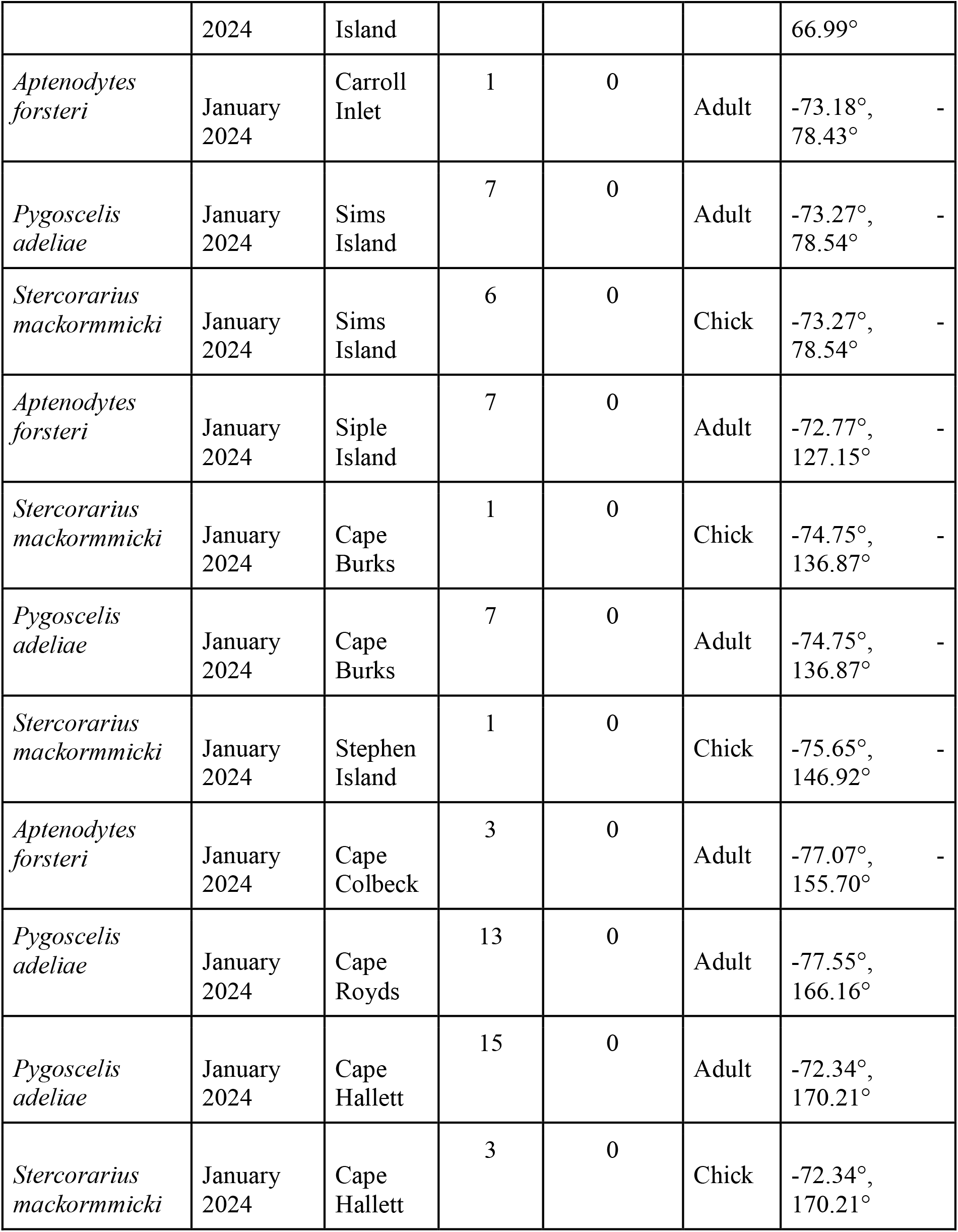
Detailed breakdown of the total number and suspected samples per species, status, period, and location sampled during the summer 2023/2024.^*^ stands for suspected cases.

**Figure 1.**
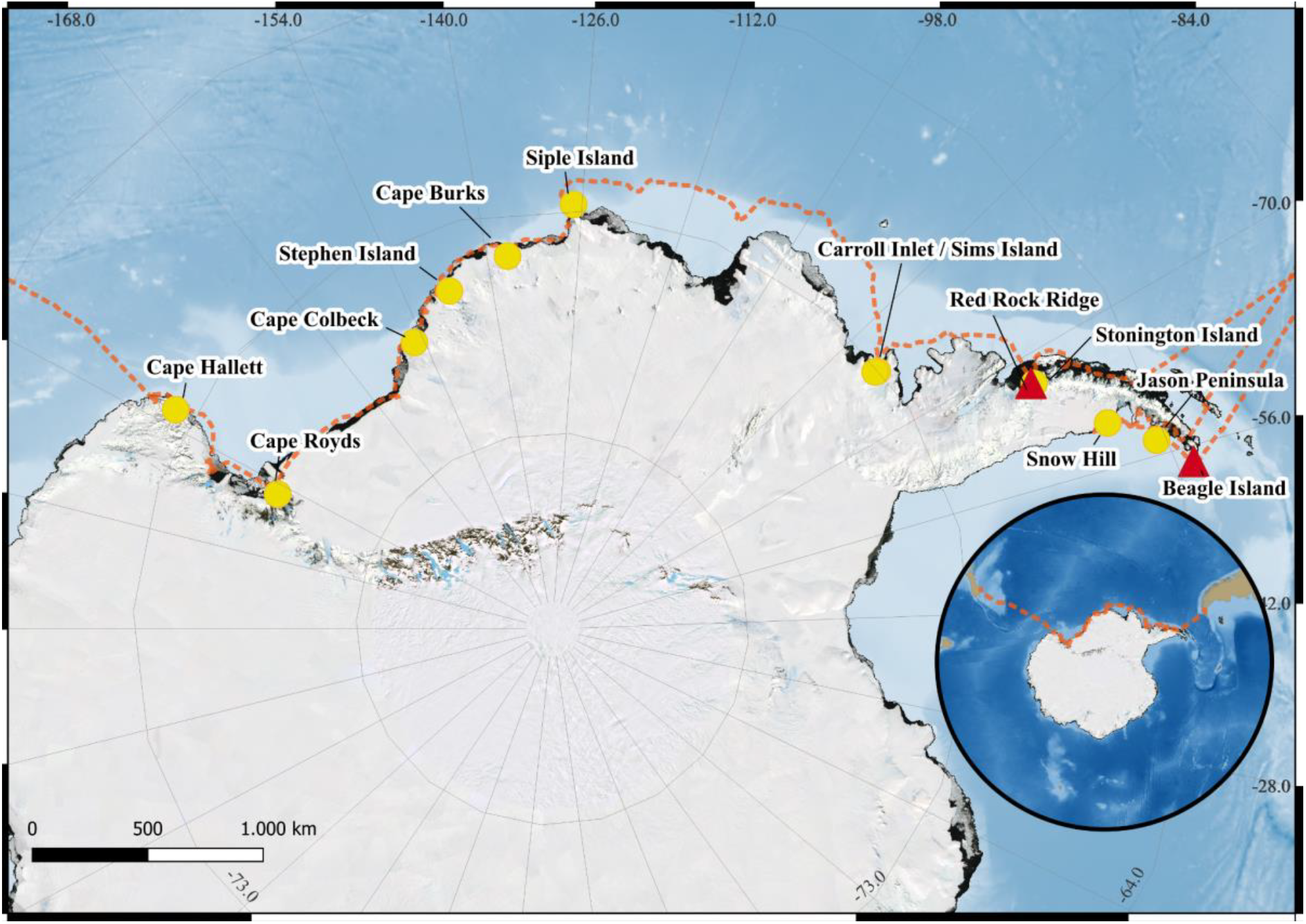
Sampling sites. Positive cases are denoted in red triangles and negative cases in yellow circles. The orange dashed line shows the ship track followed.

RNA was extracted using HiBind RNA Minicolumns (RNA extraction Omega) and amplified by RT-PCR against segment 4 of HPAI H5N1, with specific primers AH5-918F (5’-CCARTRGGKGCKATAAATC-3’) and AH5-R1618 (5’-CCARTRGGKGCKATTAATC-3’), with an expected amplicon size of 782 bp. RT-PCR conditions were retrotranscription at 45ºC for 30’, followed by denaturation of 95ºC for 3’, and 40 cycles of 95ºC for 15’’, 52ºC for 30’’, and 72ºC for 60’’, followed by final extension of 72ºC for 10’. The procedure was repeated and confirmed three times for the suspected cases and randomly chosen negative cases, with a blank control included for each PCR run.

Importantly, the 16 Adélie penguins sampled in Beagle Island were also equipped with Argos satellite trackers within the framework of another study (Le Bohec, *unpublished data*), which allowed us to assess their fate in the period following sampling.

## Results

Of the 115 individuals sampled, 9 tested positive, including 8 Adélie penguins and 1 Antarctic shag, corresponding to a positivity rate of ∼12% for all the Adélie penguins sampled. On Beagle Island (Danger Islands, at the northern tip of the Antarctic Peninsula), nearly half the Adélie penguins sampled were positive for RT-PCR amplification. All suspected positive cases came from the Antarctic Peninsula, and no suspected cases were detected in the 64 samples collected in the western part of the Bellingshausen Sea, in the Amundsen Sea, or in the Ross Sea. None of the birds sampled, including the suspect cases, showed any symptoms. Crucially, the 7 Adélie penguins that tested positive in Beagle Island and also carried an Argos tracker were still alive and engaged in breeding-like foraging behavior as of early March 2024, over 2 months after the positive test, thus establishing the low-pathogenicity of H5N1 infection in these 7 cases. Molecular analysis confirmed the presence of PCR amplicons of the expected size for H5N1 in positive cases and their replicates, while replicates of negative cases consistently yielded negative results.

## Discussion

The first-time detection of suspected cases of H5N1 influenza in penguins and cormorants in the Antarctic continent marks a significant expansion of the panzootic into this isolated continent and puts highly vulnerable bird populations at risk. Individual cases were already reported for migratory Antarctica skuas in the peninsula region in the previous months (https://scar.org/library-data/avian-flu), confirming the fear that this widely-dispersing bird may be the first vector or the virus to the continent. Our findings now establish the transmission of the virus to two Antarctic resident species. Although this study covered a large part of the West Antarctic, at this stage positive cases seem restricted to the Antarctic peninsula (Beagle Island and Red Rock Ridge in our sampling). However, it is now clear that the HPAI virus has penetrated the Antarctic continent and has a high prevalence in two highly abundant resident species. There is thus significant apprehension regarding the further spread of the virus, due to the high density and mobility of these birds and their gregarious reproductive behavior - two traits that enhance intra and inter-colony virus spread, and that are thought to have largely worsened the HPAI panzootic in Northern Hemisphere seabirds (Paradell et al. 2023)

The absence of clinical signs in potentially HPAIV H5N1-infected birds, coupled with the lack of excess mortality at the sampled sites despite a positivity rate of roughly 50% among Adelie penguins in Beagle Island, raises the possibility that some seabird species may be more susceptible to H5N1 than others, as previously reported by Chilean authorities in Humboldt and Magellanic penguins (SERNAPESCA, https://www.sernapesca.cl/influenza-aviar/). These asymptomatic cases may seem reassuring for the species in question, but they have strong implications for Antarctic wildlife in general, potentially leading to unnoticed and widespread virus transmission, as asymptomatic carriers could serve as “trojan horses,” facilitating the introduction and spread of HPAIV to previously unaffected populations, eventually contaminating more susceptible species (such as pinnipeds, who died in large numbers of HPAI in South America and South Georgia). These new findings emphasize the pressing need to monitor and address emerging diseases in vulnerable Antarctic seabirds. Implementing rigorous biosecurity measures during Antarctic activities is essential to minimize the potential spread of avian influenza.

## Acknowledgments

This study was funded by ICM-ANID ICN2021_002 Millennium Institute BASE, the Oceanographic Institute, Foundation Albert I, Prince of Monaco (IOM), the Centre Scientifique de Mónaco, ICN2021_044 – CGR. We are very grateful to the all team of ICM-ANID ICN2021_002 Millennium Institute BASE and of IOM, with the very strong support of Robert Calcagno, Cyril Gomez, Lara Bleu, Flavia Ferrucci. We deeply thank the Ponant science team and the Commandant Charcot crew, and in particular Eric Dupont, Jean-Phillippe Savy, Patrick Marchesseau, Wassim Daoud; the Sedna team, and in particular Nicolas Dubreuil; Jean-Yves Barnagaud, Paul Dufour, Aude Boutet and Elsa Freschet for their help on the field. All procedures were approved by the Terres Australes et Antarctiques Françaises (TAAF; permit #2023-184) and Instituto Antártico Chileno (INACH; sampling permits 1008/2023 and 28/2024, permits for access into protected areas 168-2024, 150-2024 and 151-2024).

## Notes

### Competing Interest Statement

The authors have declared no competing interest.

